# fsbrain: an R package for the visualization of structural neuroimaging data

**DOI:** 10.1101/2020.09.18.302935

**Authors:** Tim Schäfer, Christine Ecker

**Affiliations:** Department of Child and Adolescent Psychiatry, Psychosomatics and Psychotherapy, University Hospital Frankfurt, Goethe University Frankfurt am Main, Germany; Brain Imaging Center, Goethe University Frankfurt am Main, Germany; Department of Forensic and Neurodevelopmental Sciences, Institute of Psychiatry, Psychology and Neuroscience, King’s College London, London, United Kingdom

## Abstract

**Summary:** We introduce fsbrain, an R package for the visualization of neuroimaging data. The package can be used to visualize vertex-wise and region-wise morphometry data, parcellations, labels and statistical results on brain surfaces in three dimensions (3D). Voxel data can be displayed in lightbox mode. The fsbrain package offers various customization options and produces publication quality plots which can be displayed interactively, saved as bitmap images, or integrated into R notebooks.

**Availability and Implementation:** The software, source code and documentation are available under the MIT license at https://github.com/dfsp-spirit/fsbrain. Releases can be installed directly from the Comprehensive R Archive Network (CRAN).

## 1 Introduction

In surface-based structural neuroimaging, raw data and statistical results are typically presented on brain surface meshes. Standard software packages like FreeSurfer [Fischl(2012)] can be used to reconstruct the human cortex from three-dimensional magnetic resonance imaging (MRI) data and compute vertex-wise measures like cortical thickness based on the resulting cortical meshes. The measures from different individuals can then be mapped to a common or standard space template for group comparisons and other statistical analyses. While standard approaches like the general linear model are available in most neuroimaging software packages, these packages do not offer the flexibility and functionality of dedicated statistical software environments such as R [R Core Team(2017)]. R provides seamless access to a wide range of state-of-the-art statistical methods via a package system. However, visualization options for neuroimaging data in R remain limited, and due to the spatial (i.e. 3D) nature of surface-based data, their representation with standard 2D visualization tools is not feasible.

The cerebroViz package [Bahl, Koomar, and Michaelson(2017)] can be used to plot spatiotemporal gene expression or other data in certain pre-defined brain regions. The output is a 2D scalable vector graphics (SVG) object, in which the expression level is color-coded. The ggseg package for brain segmentations [Mowinckel and Vidal-Piñeiro(2019)] provides an application programming interface (API) inspired by ggplot2. It can be used to visualize pre-defined brain parcellations from different atlases, or region-based data on an inflated mesh. However, to the best of our knowledge, no specialized R package yet exists that supports visualizations of vertex-wise data such as derived by FreeSurfer. Here, we present the fsbrain package for R, which provides publication quality visualization of structural neuroimaging data in R. For surface-based workflows, both vertex-wise and region-based data are supported, and high-level functions exist to plot raw morphometry data in native and standard space, surface labels, brain surface parcellations, and results like statistical maps or clusters. Volume-based data (voxels) can be displayed in 3D and in lightbox view.

## 2 Implementation

The fsbrain package reads neuroimaging data and cortical meshes in different file formats used by popular neuroimaging software packages using the R packages freesurferformats [Schäfer(2020)], oro.nifti [Whitcher, Schmid, and Thornton(2011)] and gifti [Muschelli(2019)]. Data computed in R can also be plotted directly. The visualization works by (1) transforming region- or vertex-wise numerical data into vertex colors using a color-map or a lookup table, (2) rendering the vertex-colored meshes in rgl [Adler, Murdoch, and others(2020)] from different viewing angles, (3) creating a color legend, and (4) combining it with the individually rendered views into an output image. The result can be displayed interactively, saved as a bitmap image, or integrated into an R notebook.

The API of fsbrain is split into low-level functions, high-level functions, and export functions. The low-level functions allow full control over the rendering process, while the high-level functions support rendering most types of neuroimaging data in a single line of R code. Both are suitable for the live inspection of data before and during the analysis, which is crucial for evaluating analysis progress and interpreting results. The export functions serve to render publication-ready high-quality images in a tight layout. They can also be used to combine several types of figures, e.g. plots from different view angles, participants, surfaces, or features into a single final image.

The plots produced by fsbrain can be customized with different colormaps (divergent and sequential), view angles, and rendering settings like material properties.

## 3 Application

In this section, we showcase different visualization options. Figure 1a shows vertex-wise morphometry data for a single subject in standard space. Here, sulcal depth values computed in FreeSurfer were mapped to colors using the viridis colormap. In Figure 1b, the regions from the Desikan-Killiany atlas [Desikan, Ségonne, Fischl, Quinn, Dickerson, Blacker, Buckner, Dale, Maguire, Hyman, Albert, and Killiany(2006)] were loaded for an individual in native space. The average cortical thickness per region was computed and visualized on the native space surface of the participant. Figure 1c shows a *t* statistical map. The medial wall is displayed in white. Figure 1d shows clusters of significant *t* values as they are typically visualized in neuroimaging studies. A divergent colormap is used to clearly separate positive and negative *t*-values on an inflated brain surface. The curvature of the original surface is displayed in the background to indicate the positions of the clusters in relation to the gyri and sulci of the brain, which makes orientation on the inflated surface easier. An example lightbox visualization of a brain volume can be found in Figure S1 in the supplement, along with the source code used to create all figures in this article.

**Figure 1.**
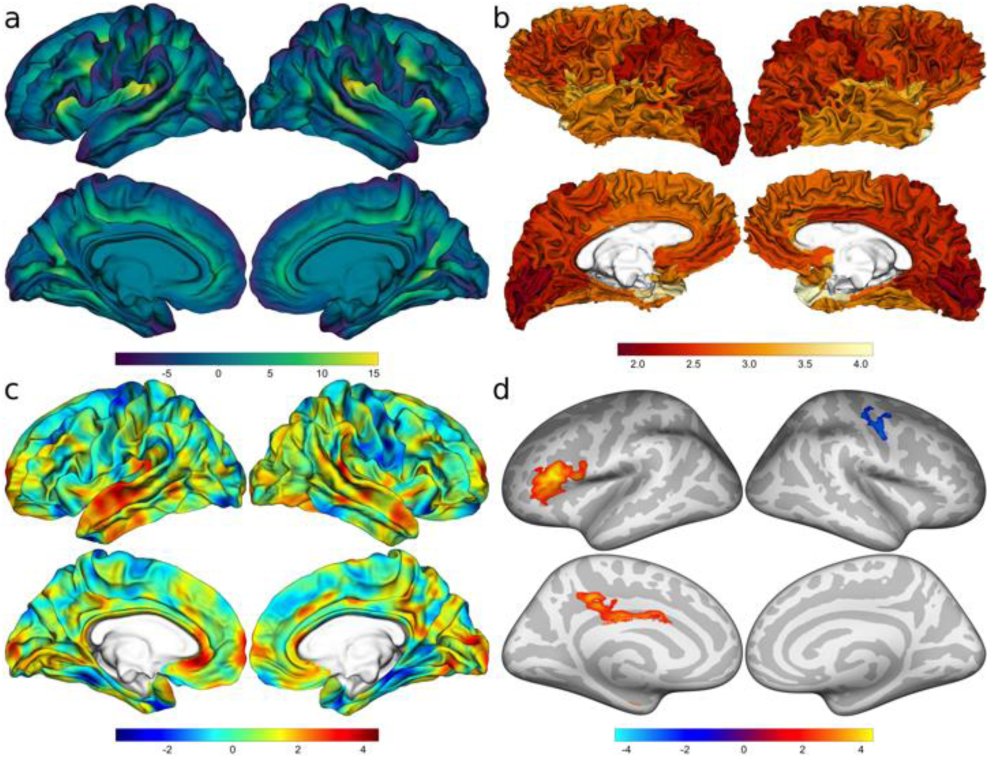
(a) Sulcal depth for a single subject. (b) Mean cortical thickness for the regions of the Desikan atlas. (c) A t value map on the fsaverage white surface. (d) Clusters on the inflated fsaverage surface. The positions of gyri and sulci are displayed in grayscale.

## 4 Conclusion

The fsbrain package provides easy-to-use functions to visualize volume-based and surface-based neuroimaging data directly in R. The package has extensively been tested, and detailed tutorials in form of R notebooks are available at the project website. It can be installed from GitHub or directly from CRAN. In combination with the statistical methods available in R and markdown notebooks, fsbrain fills the gap between analyzing and visualizing surface-based data and allows accessing the full analytical workflow: everything from data loading to the visualization of results can be carried out in the R environment. We hope this helps researchers, and facilitates reproducible analyses of neuroimaging data in R.

## Supporting information

Supplemental Material incl. Figure S1

## References

D Adler et al. rgl: 3D Visualization Using OpenGL. 2020.

E Bahl et al. cerebroViz: an R package for anatomical visualization of spatiotemporal brain data. Bioinformatics. 33(5):762–763, March 2017. doi: 10.1093/bioinformatics/btw726.

R S Desikan et al. An automated labeling system for subdividing the human cerebral cortex on MRI scans into gyral based regions of interest. NeuroImage. 31(3):968–980, July 2006. doi: 10.1016/J.NEUROIMAGE.2006.01.021.

B Fischl. FreeSurfer. NeuroImage. 62(2):774–781, August 2012. doi: 10.1016/j.neuroimage.2012.01.021.

A M Mowinckel and D Vidal-Piñeiro. Visualisation of Brain Statistics with R-packages ggseg and ggseg3d. 191208200 Stat. December 2019.

J Muschelli. gifti: Reads in “Neuroimaging” “GIFTI” Files with Geometry Information. 2019.

R Core Team. R: A Language and Environment for Statistical Computing. R Foundation for Statistical Computing, Vienna, Austria, 2017.

T Schäfer. freesurferformats: Read and Write “FreeSurfer” Neuroimaging File Formats. 2020.

B Whitcher et al. Working with the DICOM and NIfTI Data Standards in R. J. Stat. Softw. (6), October 2011. doi: 10.18637/jss.v044.i06.

