## Supplemental Material incl. Figure S1 for "fsbrain: an R package for the visualization of structural neuroimaging data": fsbrain_bioinf_supplement.html

fsbrain Supplementary Material


### fsbrain Supplementary Material

###### Tim Schäfer

###### September 7, 2020

This file contains Figure S1 and the source code to reproduce all figures included in the publication. This file was generated with fsbrain version 0.4.0.

Note that the fsaverage data is part of FreeSurfer and distributed under the terms of the FreeSurfer software license.

##### Setup

Here we download required data and set the subjects directory.

```
library('fsbrain');
fsbrain::download_optional_data();
fsbrain::download_optional_paper_data();
fsbrain::download_fsaverage(accept_freesurfer_license = TRUE);
sjd = fsbrain::get_optional_data_filepath('subjects_dir');
sj = 'subject1';
fsbrain.set.default.figsize(1000, 1000);
```

#### Reproduce Figure 1

##### Figure 1a (standard space morphometry data: sulcal depth)

```
cm1a = vis.subject.morph.standard(sjd, sj, 'sulc', rglactions=list('no_vis'=T));
fig1a = vis.export.from.coloredmeshes(cm1a, colorbar_legend='Sulcal depth [mm]');
```

The image gets exported as a PNG file into the current working directory by default, and you can set the ‘output\_img’ parameter to save it wherever you want. One can also show it directly:

```
fig1a;
```

##### Figure 1b (region-based data)

```
lta = group.agg.atlas.native(sjd, sj, 'thickness', 'lh', 'aparc');
rta = group.agg.atlas.native(sjd, sj, 'thickness', 'rh', 'aparc');
cm1bt = vis.region.values.on.subject(sjd, sj, 'aparc', lta, rta);
fig1b = vis.export.from.coloredmeshes(cm1bt);
```

```
fig1b
```

##### Figure 1c (t map)

```
tmaps = subject.morph.standard(sjd, sj, 'tmap', hemi='both', fwhm=NULL, cortex_only = T, split_by_hemi = T);

cm1c = vis.data.on.fsaverage(morph_data_lh = tmaps$lh, morph_data_rh = tmaps$rh, makecmap_options = list('colFn'=squash::jet, 'n'=500, col.na = 'white'));
fig1c = vis.export.from.coloredmeshes(cm1c);
```

```
fig1c;
```

##### Figure 1d (clusters)

```
lh_demo_cluster_file = system.file("extdata", "lh.clusters_fsaverage.mgz", package = "fsbrain", mustWork = TRUE);
rh_demo_cluster_file = system.file("extdata", "rh.clusters_fsaverage.mgz", package = "fsbrain", mustWork = TRUE);

lh_clust = freesurferformats::read.fs.morph(lh_demo_cluster_file);
rh_clust = freesurferformats::read.fs.morph(rh_demo_cluster_file);

cm1d = vis.symmetric.data.on.subject(sjd, 'fsaverage', lh_clust, rh_clust, bg="curv_light", surface = 'inflated');
cm1d = shift.hemis.apart(cm1d);
fig1d = vis.export.from.coloredmeshes(cm1d);
```

```
fig1d;
```

#### Create Figure S1: lightbox

```
brain = subject.volume(sjd, sj, 'brain');
aparc_aseg = subject.volume(sjd, sj, 'aparc+aseg');

colortable = freesurferformats::read.fs.colortable(file.path(sjd, 'fsaverage', 'ext', 'FreeSurferColorLUT.txt'));
overlay_colors = vol.overlay.colors.from.colortable(aparc_aseg, colortable);
colored_brain = vol.merge(brain/max(brain), overlay_colors);

figS1 = volvis.lightbox(colored_brain);
```

```
figS1;
```

Figure S1: Lightbox view of a brain volume. The original volume has dimensions 256x256x256. Here, starting from the first non-empty slice, every fifth slice along one axis is visualized. The results of the FreeSurfer ‘aparc+aseg’ segmentation are displayed using the colors of the FreeSurfer color table. The original brain image is displayed in the background in grayscale.
